# Reliability metrics of outcome measures during the single and double leg drop jump tests

**DOI:** 10.1101/2022.07.10.499453

**Authors:** Huseyin Celik, Ahmet Yildirim, Evrim Unver, Caner Mavili, Ekrem Yilmaz, Ferhat Ozturk, Pinar Arpinar-Avsar, Sukru Alpan Cinemre

## Abstract

Although a number of previous studies have studied reliability metrics of outcome measures during single and double leg drop jump tests, none of them presented detailed reliability information for both tests with a group of collegiate athletes of both genders. The aim of this study was therefore to assess the reliability of outcome measures during the single and double leg drop jump tests with proper reliability metrics. Seventeen handball players (11 male and 6 female) participated in the experiments. Each player performed three double leg and three single drop jumps from a 30 cm height box onto a portable force plate in two sessions a week apart. Sixteen outcome measures, including jump height (JH), ground contact time (GCT), reactive strength index (RSI), were calculated. Four groups of reliability metrics like absolute agreement and consistency intraclass correlation coefficients (inter-day and intra-day), standard error of measurement, coefficient of variation, and minimal metrically detectable change were estimated for reliability assessment. The outcome measures as RSI and its individual components JH and GCT together with normalized vertical stiffness (Kvert) yielded high inter-day and intra-day intraclass correlation coefficients and low standard error of measurement and coefficient of variation levels for the double leg drop jump test. The more challenging single leg drop jump test could also be considered reproducible with some other highly reliable outcome measure set by keeping JH, GCT, and normalized Kvert and replacing vertical jump impulse with RSI. The results of the current study therefore suggested the drop jump tests could be deemed reliable to be used for short-term and long-term monitoring needs for a group of collegiate athletes of both genders.

## Introduction

Muscle exercises are classified primarily into static and dynamic types (Knuttgen ve Komi, 2003). Plyometrics is a dynamic type of exercise included in training programs to improve speed, endurance, reactive strength, and explosive power of athletes (e.g., Chu and Myer, 2013; Slimani et al., 2016). Plyometric exercise comprises an eccentric stretch of the muscle tendon unit (MTU) immediately followed by a concentric shortening of the MTU. This combination of MTU undergoing active stretching followed by rapid shortening during the stretch-shortening cycle (SSC) is the foundation of plyometric exercise which seeks to exploit the force potentiating capabilities of the SSC (Norman and Komi, 1979). Different types of jumps such as tuck jump, single leg hop, and depth or drop jump are commonly used in plyometric exercise programs to improve vertical jumping ability (Chu and Meyer, 2013).

One of the most popular vertical jump drills in plyometric training programs is drop jump (DJ) (Bobbert, 1990). DJ consists of standing on a box, stepping off and dropping onto the ground, upon landing (eccentric stretching or braking phase), immediately performing a vertical jump (concentric shortening or propulsion phase) (Komi, 2003). One of the proposed mechanisms responsible for the force potentiation observed in SSC is the recovery of elastic potential energy (e.g. Alexander, 1987; Blazevich, 2011). The elastic potential energy is stored in the MTU during the eccentric stretch phase and that stored elastic energy is released during the concentric shortening phase, and MTU loading is the dominant factor that influences energy storage (Blazevich, 2011). Baxter and his colleagues (2021) evaluated Achilles tendon loading profiles of various exercises and proposed that single and double leg drop jumps are tier 3 and 4 (the highest tier) respectively. This research indicated that single and double leg drop jumps could be assessed separately as loading profiles are different, yet little research has been done on single leg DJ compared to double leg DJ.

Several previous studies examined double leg DJ and compared it with other vertical jumps using biomechanical instrumentation such as electromyography systems, motion capture cameras, and force plates (e.g. Bobbert et al., 1987; Ishikawa et al., 2006; Healy et al., 2014). In one of the first studies on the subject, Komi and Bosco (1978) studied the vertical jump performance of men and women for three types of jumps as squat jump, countermovement jump, and drop jump. In the many following studies, vertical jump performance has been studied extensively from the point view of biomechanical performance, neuromuscular control, and SSC function variables using total body biomechanics (e.g. Harry et al., 2021; McMahon et al., 2021). In order to do so, several outcome measures, such as jump height, ground contact time, reactive strength index have been calculated from the force-time curve variables recorded by a force plate during vertical jumps (McMahon et al., 2021). Those outcome measures have the potential to be useful for profiling and benchmarking, injury prevention, and rehabilitation and return to sport perspectives, and to be used to evaluate effectiveness and impact of vertical jump drills in plyometric training programs, if obtained in a reproducible way.

To be capable of identifying meaningful change for both short-term and long-term monitoring needs, reliable and repeatable test protocols are critical to creating a performance evaluation system (French et al., 2022). Sport scientists who evaluate performance have to make sure that the procedures they follow do not involve errors that are larger than the differences they are supposed to assess and also the outcome measures are reliable to count them on (Kibele, 1998). Thus, the jump data obtained from tests must be reproducible and consistent, in other words, the following measurements under the very same conditions should yield virtually the same results as the first measurement. This reproducibility is usually quantified using reliability metrics by implementing a test–retest method in which the first data set obtained in the first measurements is compared with the data obtained from the same subjects in the following measurements at some other time under the very same conditions (Vincent and Weir, 2020). Such reliability metrics as intraclass correlation coefficient (ICC) and standard error of measurement are reported frequently to quantify test-retest reliability in the exercise and sport sciences (Weir, 2005).

The reliability evidence, however, often lacks clarity (Morrow and Jackson, 1993). The methodology used to obtain outcome measures and reliability metrics might not be defined in detail in the published studies (Liljequist et al., 2019). Also, studies with the purpose of establishing the reliability of tests should be representative of the defined population in terms of age and gender (Morrow and Jackson, 1993). However, as noted in a very recent study (Harry et al., 2021), most of the generated data on vertical jumps were stemming from male-only studies. Actually, this situation is not limited to studies on jumping, since it has been demonstrated quantitatively in the article “Invisible Sportswomen”: The Sex Data Gap in Sport and Exercise Science Research (Cowley et al., 2021). The authors examined the gender distribution of approximately 12 million participants in 5261 articles published in the field of sports and exercise sciences, 63% of the publications presented data on both male and female participants, 31% were only men, and 6% only women (approximately 8.2 million male and 4.2 million female participants in total). As the results of the study (Cowley et al., 2021) clearly demonstrated, women are significantly underrepresented in sport and exercise science research.

Despite the scarcity of data on woman, scientific papers, technical notes, and new technological improvements are piling up about measurement, training, and analysis methods for vertical jumps (e.g. Comfort et al., 2019). With new technological improvements, portable instruments such as portable force plates have been started to be used in jump tests with a field test approach (e.g. Uzelac-Sciran et al., 2020). There is however limited research investigating the reliability of outcome measures of DJ tests measured with portable force plates, even less for single-leg DJs. Previous research mostly focused on double leg DJs measured with a stationary force plate from male subjects. For instance, Flanagan and his colleagues (2008) used a stationary force plate and examined the intra-day reliability of the outcome measures of the double leg DJ test. The authors found that jump height, ground contact time, and reactive strength index were to be highly reliable from trial to trial evidenced by very high ICC values. Conversely, TTS was deemed not reliable from trial to trial, as evidenced by moderate to low ICC values. However, arm position was not controlled throughout the jumping and landing movements which could be a reason for that result. Also, the gender of the subjects was not specified explicitly despite the fact that research on vertical jumps have suggested there are several differences between male and female subjects (e.g. Komi and Bosco, 1978; Laffaye et al., 2010). In a more recent study, Byrne and his colleagues (2017) investigated the inter-day reliability of the reactive strength index and optimal drop height using an optical system for measurements. The subjects were 19 male hurling players. The reactive strength index and optimal drop height measures yielded high reliability with ICC values of 0.87 and 0.80 as well as coefficient of variation values of 4.20% and 2.98% respectively. However, this study was also performed with only male subjects. In addition to that an optical not a force plate system was used for the measurements. Moreover, the details of ICC computations and models were not reported in detail like many other studies (Liljequist et al., 2019).

As vertical jumping is regarded as a central and fundamental element of many sports, vertical jumping ability is one of the factors that determine performance in a variety of sports (Bobbert, 1990; Aragon-Vargas, 2000). Handball, a contact team sport which includes high-paced and short-term activities such as jumping, is not an exception to that. Due to the nature of handball, the players’ biomechanical performance, neuromuscular control, and SSC function were emphasized. Therefore, monitoring variables related with those aspects by means of outcome measures of DJ tests might be beneficial. Given also the resemblance of single leg jumps to some fundamental movements in handball suggests that studying single leg DJ as well as double leg DJ may be worthwhile.

This experimental study will test the hypothesis that outcome measures of single and double leg DJ tests such as jump height, ground contact time, reactive strength index calculated from a group of male and female handball players using a portable force plate would yield reliable measurements from trial to trial and day to day. A repeated measures experimental design was implemented with subjects performing three single leg and three double leg DJs from a fixed height in a non-fatigued state. The very same experimental measurements were repeated after one week. The inter-day and intra-day reliability of each of the outcome measures will be statistically assessed with reliability metrics as ICC, standard error of measurement, coefficient of variation, and minimal metrically detectable change. Ee shared all data and code repository on Open Science Framework. Hence, anyone interested could be able to repeat the analysis and change any procedure in the methodology to assess possible effects of them on the reliability metrics and outcome measures.

## Materials and Methods

### Subjects

Seventeen voluntary participants (11 male and 6 female) took part in the experiments (age: 20.9±2.5 years, height: 176.8±8.3 cm, weight: 71.6±14.3 kg) with no neuromusculoskeletal problem that would affect their jumping performance at the time of measurements. The subjects were the members of the college handball team. The study took place at METU, Ankara, Turkey. All subjects have experience in plyometric exercise but did not perform such exercises at the time of the study. Subjects had performed no strength training in the 48 hours before data collection. All subjects gave informed consent to participate in the study. This specific study was reviewed and approved by Non-interventional Clinical Researches Ethics Board of Hacettepe University (No: GO 17/902).

A statistical power analysis was performed a priori for sample size estimation. With a correlation coefficient, r = 0.80 and power = 0.90, the projected sample size needed was 11 subjects (Portney and Watkins, 2015). We recruited 17 members of the college handball team. Previous studies on the outcome measures of drop jump tests indicated that this sample size would be adequate in attaining acceptable levels of power in reliability analysis studies (the number of subjects were 22, 18, and 19 at Flanagan et al. (2008), Lloyd et al. (2009), and Byrne et al. (2017) studies respectively).

### Instrumentation

During DJs, the ground reaction force in the vertical direction (VGRF) were registered at 2000 Hz with a portable single force plate (Kistler 9260AA, Switzerland)) for the sample period. The computer software interface was Kistler’s own software MARS. Each raw data of each trial was stored on the same Pentium-powered laptop and then exported as comma-separated values (csv) file to MATLAB environment for calculations of outcome measures and reliability metrics.

### Procedures

Before testing, a week earlier, the athletes completed a familiarization session in which they had opportunity to practice single and double DJs. Then, the recorded DJ tests were administered by the same raters in two sessions a week apart at the same time of day, at the same indoor venue where the collegiate athletes train, on a level sports court with a synthetic hard floor. Before each measurement session, the athletes were re-informed about the procedures. The participants wore the same shoes and similar clothing for the tests, and were to avoid drinking (other than water), eating, and exercising in the hour before testing. DJ tests were assessed in two different conditions as double and single leg; first three trials of double leg, then three trials of single leg to provide the participants with a gradual increase in neuromuscular stress (Lloyd et al., 2009).

Before the measurements, the subjects were then given verbal description and visual demo of single and double leg DJs. The instruction set was: (i) step forward off the box of 30 cm without stepping down and drop onto the force plate, (ii) upon landing perform a rebound jump as high as possible and as quickly as possible, think the force plate as a hot plate (iii) on touch-down stick your landing and to stabilize as quickly as possible while facing straight ahead and remain motionless by simply looking at a cross sign on the wall approximately 4 m in front of you, (iv) keep your hands on the hips throughout the jump, (v) do not tuck your legs in the air (Dalleau et al., 2004; Flanagan et al., 2008; Byrne et al., 2017). In both DJ tests, the participants stepped forward off the box with their preferred foot which was established in the familiarization session and remained the same in the following sessions (Maloney et al., 2016).

The warm-up procedure consisted of five minutes of jogging, followed by dynamic stretching of each major muscle group of the lower limbs (Flanagan et al., 2008). After warm-up, the participants were provided with the opportunity to practice both jumps. Then the participants were allowed for five minutes of rest before the DJ tests. The participants performed six maximal DJs (three double and three single leg). The rest period was approximately one minute between the jumps. If the participant could not stabilize in single leg DJ, that trial was repeated. Data was successfully registered for all participants for both jumps.

### Data Analysis

The data analysis had two stages: (i) detection of key time points such as landing and touch-down instants, (ii) calculation of outcome measures of DJs. Both single and double leg DJ performance of the participants were analyzed with the exact same code based on the methods described by Baca (1999). Fig 1 showed an example of VGRF record for a double leg DJ. Fig 1 also indicated key time points of the jump. We used the same numbering order as in Baca (1999). Specifically, t2 was identified as the instant of landing onto the force plate by locating the data point where VGRF value first exceed 10 N threshold (Lloyd et al., 2009; Castilla et al., 2021) after starting recording.

**Fig 1.**
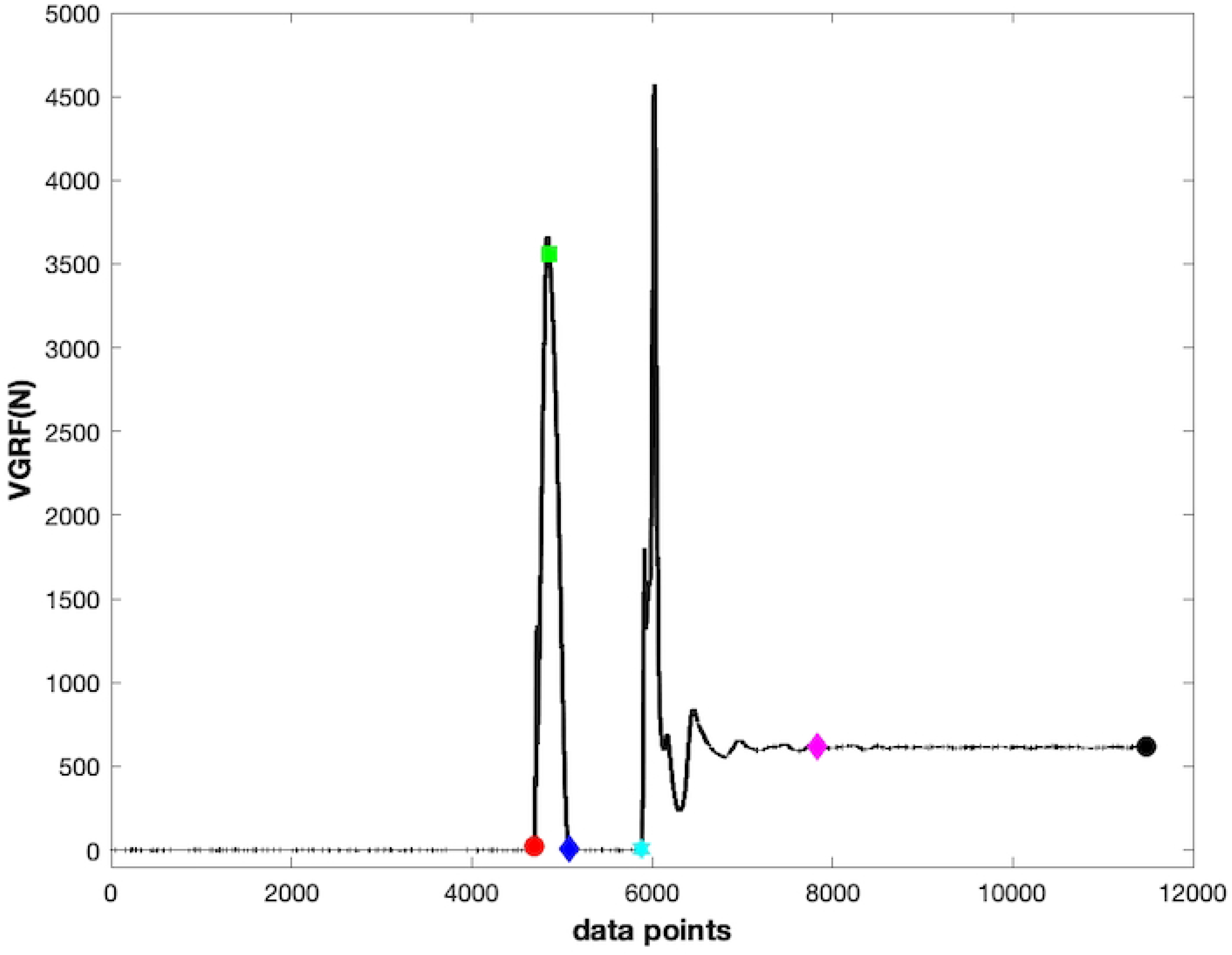
Exemplary force-time curve of the vertical ground reaction force (VGRF) during a double leg drop jump. t2 (red circle): landing onto the portable force plate, t3 (green square): the instant where vertical velocity of total body center of mass is zero, t4 (blue diamond): the instant of take-off, t5 (cyan hexagram): the instant of touch-down, t6 (black circle): the last point of force record at which the participant stands still on the force plate, vertical time to stabilization (magenta diamond): defined below, from t2 to t3 is the eccentric or braking phase, from t3 to t4 concentric or propulsion phase.

Similarly, t5 was identified as the instant of touch-down on the force plate by locating the data point where VGRF value first exceed 10 N threshold after the maximal vertical jump. The take-off instant, t4, was located as the data point where the VGRF value first drops below 10 N threshold after the t2 instant. The t6 is the last point of force record at which the participant stands still on the force plate. The vertical velocity of total body center of mass (BCOM) at the instant of landing (t2) was estimated by numerically integrating the vertical force minus body weight (BW = m*g where m is the body mass and g is the gravitational acceleration, 9.80665 m/s^2^) from t6 to t2 using the trapezoid method (Baca, 1999).

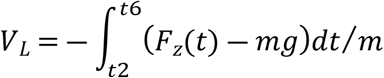

The t3 is the instant where the vertical velocity of BCOM is zero, and located with solving the following equation below numerically (Baca, 1999).

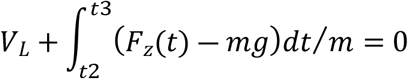

The t3 instant divides ground contact phase (from t2 to t4) into two as eccentric or braking phase (from t2 to t3) and concentric or propulsion phase (from t3 to t4) (Baca 1999; Komi 2003). The t1 instant was when participant take-off from the box, yet we did not locate that instant as Baca (1999) used a second force plate for that purpose.

After locating the key time points, we calculated several outcome measures of DJs. Kinetic analysis of vertical jumping by using force plates enables extensive assessment of neuromuscular function in eccentric (braking) and concentric (propulsion) phases of SSC separately and jointly, allowing researcher and practitioners to go beyond testing only simple measure of jump height (Bishop et al., 2021). Recently, Bishop and his colleagues (2021) published a framework for selecting outcome measures during DJ tests. The authors suggested reactive strength index and its individual components jump height and ground contact time as well as lower extremity stiffness. Other outcomes measures as time to stabilization, peak maximal VGRF, peak BCOM displacement, rate of force development, mean braking and propulsion force, braking and propulsion phase time were also reported in the previous studies (Flanagan et al., 2008; Maloney et al., 2017; Sarabon et al., 2020; McMahon et al., 2021). Besides, in his recent book, Cleather (2021) argued that impulse would be the most important outcome measure in explosive physical performances. Therefore, we have included impulse measures to our analysis.

From the force-time curves obtained from the force plate, we calculated the following outcome measures.

- **Jump height** (JH in cm): JH was the peak displacement attained by the BCOM during the flight phase (from t4 to t5 in Fig 1) and calculated using the flight time in the air (JH-TIA) and the vertical velocity of BCOM the take-off instant (JH-TOV) (Linthorne 2001; Moir, 2008).
- **Ground contact time** (GCT in s): GCT was the time spent on the ground after landing until take-off (from t2 to t4 in Fig 1) and used to classify SSC as fast (100-250 ms) and slow (>250 ms) (Schmidtbleicher, 1992).
- **Reactive strength index** (RSI in cm/s): RSI has been used to quantify SSC performance and calculated as the ratio of JH to GCT (Flanagan and Comyns, 2008).
- **Braking phase time** (BPT in s): BPT was the time spent on the ground in the eccentric braking phase and calculated as the time between the landing instant and the following first instant of zero vertical velocity of BCOM (from t2 to t3 in Fig 1) (McMahon et al., 2021).
- **Propulsion phase time** (PPT in s): PPT was the time spent on the ground in the concentric propulsion phase and calculated as the time between the first instant of zero vertical velocity of BCOM after landing and take-off (from t3 to t4 in Fig 1) (McMahon et al., 2021).
- **Mean braking force** (MBF in BW): Being able to tolerate high braking forces were linked with reactive strength and calculated as the average of each vertical force data point in the braking phase (from t2 to t3 in Fig 1) (McMahon et al., 2021).
- **Mean propulsion force** (MPF in BW): Being able to exert high propulsion forces were linked with vertical jump performance (Garhammer and Gregor, 1992) and calculated as the average of each vertical force data point in the propulsion phase (from t3 to t4 in Fig 1) (McMahon et al., 2021).
- **Peak maximal ground reaction force** (PGRF in BW): PGRF was the largest VGRF encountered during the jump and calculated as the global maximum value of the VGRF data points in the contact phase (braking + propulsion, from t2 to t4 in Fig 1) (Lloyd et al., 2009).
- **Peak center of mass displacement** (PCOMD in cm): PCOMD was the largest vertical displacement during the jump and calculated as the maximum displacement value of the vertical displacement data points in the contact phase (from t2 to t4 in Fig 1) after the instant of landing onto the force plate. The vertical displacement during the contact phase was calculated from the double integration of vertical acceleration of BCOM with respect to time (from t2 to t4 in Fig 1) with integration constants for velocity as *V*(*t*=t2) = *V_L_* and for position as *Y*(*t*=t2) = 0 (arbitrarily) (Baca, 1999).
- **Rate force development** (RFD in BW/s): RFD has been used to assess explosive strength of athletes and proposed to be determined by the capacity to exert maximal voluntary effort in the early phase of an explosive movement (Maffiuletti et al., 2016). RFD was calculated as the peak value of the time derivate of the filtered vertical force-time curve in the braking phase (from t2 to t3 in Fig 1). The vertical force signal was only filtered for RFD analysis using a second order low-pass zero-lag Butterworth filter with a 5 Hz cut-off frequency (Sarabon et al., 2020).
- **Vertical jump impulse** (VJI in BW*s): The vertical impulse applied during the jump is equal to the momentum change (momentum of an object is equal to the mass of the object times the velocity of it). The impulse-momentum relation requires that impulse or total VGRF applied to the BCOM is directly proportional to the change in the vertical velocity (Cleather, 2021). VJI has been used to assess performance (Cleather, 2021) and deficits before return to sport after injury (Costley et al., 2021). VJI was calculated by integrating the vertical force-time curve during the contact phase of jump (from t2 to t4 in Fig 1) and dividing the result by the BW.
- **Absolute vertical stiffness** (Absolute Kvert in kN/m): Absolute Kvert describes the vertical displacement of the BCOM in response to VGRF during a movement (Latash and Zatsiorsky, 1993) and proposed as a representative outcome measure of summative stiffness of lower limbs (Maloney and Fletcher, 2021). Absolute Kvert was calculated as the ratio of the peak VGRF in the contact phase of the jump (from t2 to t4 in Fig 1) and displacement of the BCOM values (McMahon and Cheng, 1990).
- **Normalized vertical stiffness** (Normalized Kvert in N/m/kg): Normalized Kvert has been linked with injury risk and athletic performance (Butler et al., 2003) and was calculated by dividing the absolute Kvert value by the body mass (Maloney et al., 2016). The logic behind this normalization is that vertical stiffness has been suggested to be affected by body size (Farley et al., 1993).
- **Peak propulsion power** (PPP in W/kg): PPP was defined as the maximal power attained during the propulsion phase of the jump (Bishop et al., 2020). PPP generating capacity was related to success in explosive activities such as jumping (Haff and Stone, 2015). PPP was calculated by first obtaining the power-time curve as the dot product of vertical force and velocity, and then locating the global maximum value in the propulsion phase (from t3 to t4 in Fig 1). The PPP values were reported after normalizing by body mass.
- **Vertical time to stabilization** (VTTS in s): VTSS is an outcome measure of jumps and linked with neuromuscular control that incorporates sensory and mechanical systems in landing from a jump (Wikstrom et al., 2004; Flanagan et al., 2008). There are several methods to calculate VTTS (Fransz et al., 2015). In this particular study, a method called as sequential estimation has been used (Colby et al., 1999). Specifically, sequential average is calculated as a cumulative average of the VGRF data points after touch-down (t5 in Fig 1) by successively adding one data point at a time, hence after the first point, the average of the first 2 data points was calculated, then the average of the first 3 data points was calculated, and so on. VGRF was considered to be stable and VTSS was located when the sequential average remained within one quarter standard deviation of the VGRF in the interval three seconds after the touch-down (Colby et al., 1999).

### Statistical analysis

For the day-to-day or test-retest or inter-day reliability analysis, the mean values of three trials of the DJ outcome measures in single and double leg tests were used. We performed a paired t-test on the difference of mean values of outcome measures obtained at test and retest sessions to verify absence of systematic bias (Atkinson and Nevill, 1998). Alpha level was set at 0.05 level for all statistical analyses. For the trial-to-trial or intra-day reliability analysis, three trials of the DJ outcome measures from the retest session were used only.

To estimate relative reliability, we used two-way random effects model of ICC, which essentially compares the within-subject variability of the measurement data with the between-subject variability of the measurement data. (Shrout and Fleiss, 1979; McGraw and Wong 1996). The test-retest and trial-to-trial correlations were quantified with two forms of ICC, namely, absolute agreement ICC(A,k) and consistency ICC(C,k) (McGraw and Wong, 1996; Liljequist et al., 2019). For each ICC value, the lower and upper bounds of 95% confidence interval (CI) were also calculated by using the equations presented at McGraw and Wong (1996). The ICC is a real number usually between 0 and 1 (Liljequist et al., 2019). According to Munro’s classification, the strength of correlation coefficients was interpreted as follows to describe the degree of relative reliability: 0.00–0.25, little, if any correlation; 0.26–0.49, low correlation; 0.50–0.69, moderate correlation; 0.70–0.89, high correlation, and 0.90–1.00, very high correlation (Carter and Lubinsky, 2016).

To estimate absolute reliability, we calculated three reliability metrics. The first one was the standard error of measurement (SEM), which was calculated as the square root of the mean square error term from the repeated measures analysis of variance (Atkinson and Nevill, 1998). SEM was expressed in the actual units of outcome measures; therefore, it is practical to interpret it as the smaller the SEM the more reliable the measurements (Atkinson and Nevill, 1998). The second absolute reliability metric was minimal metrically detectable change (MMDC), the amount of change that could be considered different between two measurements and calculated as the 95% CI of SEM of the outcome measures (i.e. ±1.96 SEM) (Corriveau et al., 2000, Salavati et al., 2009). Also, the coefficient of variation (CV), basically the ratio of the standard deviation (SD) to the mean of the data, was calculated to assess absolute reliability of the outcome measures. To do so, the mean CV value from the individual CV values was computed (Atkinson and Nevill, 1998). For the trial-to-trial CV calculations, the second and third trials have been taken into calculations. The following equations were used to compute the CV metric (*n*=number of subjects, *x*_1_ and *x*_2_ are measurements for the *i*th subject (Shechtman, 2013)).

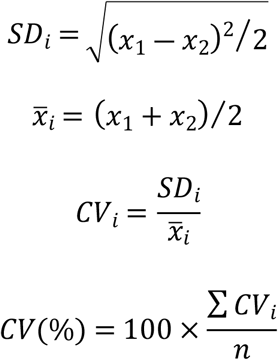

## Results

Table 1 presents the mean and SD values of 16 outcome measures of single and double leg DJ tests for test and retest measurements. Table 2 and 3 present absolute agreement and consistency ICCs and their 95% CI (in parenthesis in Table 2 and 3), SEM, MMDC, and CV metrics of the outcome measures of single and double leg DJ tests respectively.

**Table 1.** Descriptive data for outcome measures during the single and double leg drop jump tests

|  | Double leg DJ |  | Single leg DJ |  |
| --- | --- | --- | --- | --- |
|  | Test mean (SD) | Retest mean (SD) | Test Mean (SD) | Retest mean (SD) |
| JH-TIA | 21.90 (5.63) | 22.34 (6.25) | 10.09 (4.92) | 10.61 (4.17) |
| JH-TOV | 21.83 (5.71) | 22.59 (6.33) | 9.74 (5.31) | 10.21 (4.23) |
| GCT | 0.27 (0.04) | 0.27 (0.05) | 0.34 (0.05) | 0.34 (0.05) |
| RSI | 0.83 (0.28) | 0.83 (0.25) | 0.31 (0.21) | 0.31 (0.13) |
| BPT | 0.12 (0.02) | 0.13 (0.02) | 0.15 (0.02) | 0.15 (0.02) |
| PPT | 0.14 (0.02) | 0.14 (0.02) | 0.19 (0.04) | 0.19 (0.03) |
| MBF | 2.74 (0.34) | 2.74 (0.37) | 2.23 (0.27) | 2.18 (0.24) |
| MPF | 2.47 (0.36) | 2.49 (0.33) | 1.74 (0.35) | 1.76 (0.23) |
| PGRF | 4.49 (0.86) | 4.46 (0.89) | 3.69 (0.52) | 3.55 (0.50) |
| PCOMD | 18.12 (2.17) | 18.19 (2.98) | 16.83 (1.38) | 16.77 (1.83) |
| RFD | 35.49 (5.01) | 35.61 (5.38) | 29.09 (3.60) | 28.55 (3.42) |
| VJI | 0.70 (0.03) | 0.70 (0.06) | 0.66 (0.038)) | 0.66 (0.05) |
| Absolute Kvert | 23.23 (6.88) | 22.98 (8.44) | 23.26 (5.01) | 22.73 (5.47) |
| Normalized Kvert | 331.78 (100.58) | 321.57 (100.00) | 330.52 (70.67) | 324.22 (83.54) |
| PPP | 51.74 (11.69) | 52.96 (11.11) | 28.39 (11.16) | 28.22 (6.77) |
| VTTS | 1.05 (0.11) | 1.08 (0.06) | 1.12 (0.07) | 1.15 (0.11) |

**Table 2.** Reliability analysis of outcome measures during single leg drop jump test

|  | Inter-day reliability metrics |  |  |  |  | Intra-day reliability metrics |  |  |  |  |
| --- | --- | --- | --- | --- | --- | --- | --- | --- | --- | --- |
|  | ICC(A,k) | ICC(C,k) | SEM | MMDC(±) | CV% | ICC(A,k) | ICC(C,k) | SEM | MMDC(±) | CV% |
| JH-TIA | 0.75 (0.32 0.91) | 0.75 (0.31 0.91) | 2.88 | 5.65 | 17.87 | 0.97 (0.93 0.99) | 0.97 (0.94 0.99) | 1.17 | 2.30 | 11.12 |
| JH-TOV | 0.75 (0.32 0.91) | 0.75 (0.31 0.91) | 3.04 | 5.95 | 19.64 | 0.97 (0.92 0.98) | 0.97 (0.92 0.98) | 1.35 | 2.65 | 12.88 |
| GCT | 0.88 (0.68 0.95) | 0.88 (0.67 0.95) | 0.03 | 0.05 | 6.60 | 0.95 (0.88 0.98) | 0.95 (0.88 0.97) | 0.02 | 0.04 | 4.32 |
| RSI | 0.64 (0.00 0.87) | 0.63 (0.00 0.86) | 0.13 | 0.26 | 21.11 | 0.97 (0.92 0.98) | 0.97 (0.92 0.98) | 0.04 | 0.08 | 14.10 |
| BPT | 0.81 (0.47 0.93) | 0.80 (0.45 0.92) | 0.01 | 0.03 | 6.58 | 0.93 (0.84 0.97) | 0.93 (0.83 0.97) | 0.01 | 0.02 | 5.40 |
| PPT | 0.89 (0.71 0.96) | 0.89 (0.69 0.96) | 0.02 | 0.03 | 7.39 | 0.95 (0.88 0.98) | 0.95 (0.88 0.97) | 0.01 | 0.03 | 4.80 |
| MBF | 0.68 (0.14 0.88) | 0.68 (0.12 0.88) | 0.18 | 0.35 | 4.91 | 0.96 (0.91 0.98) | 0.96 (0.90 0.98) | 0.08 | 0.16 | 3.50 |
| MPF | 0.73 (0.23 0.90) | 0.72 (0.22 0.89) | 0.20 | 0.39 | 7.02 | 0.97 (0.94 0.98) | 0.97 (0.93 0.98) | 0.07 | 0.13 | 3.67 |
| PGRF | 0.45 (0.00 0.80) | 0.45 (0.00 0.80) | 0.43 | 0.84 | 8.84 | 0.91 (0.78 0.96) | 0.91 (0.80 0.96) | 0.25 | 0.50 | 6.94 |
| PCOMD | 0.70 (0.16 0.89) | 0.69 (0.15 0.89) | 1.11 | 2.18 | 5.17 | 0.78 (0.50 0.91) | 0.78 (0.49 0.91) | 1.51 | 2.96 | 5.18 |
| RFD | 0.67(0.10 0.88) | 0.67 (0.09 0.88) | 2.47 | 4.85 | 5.76 | 0.96 (0.89 0.98) | 0.96 (0.90 0.98) | 1.23 | 2.41 | 4.32 |
| VJI | 0.80 (0.45 0.93) | 0.79 (0.44 0.92) | 0.03 | 0.06 | 2.82 | 0.92 (0.82 0.96) | 0.92 (0.81 0.96) | 0.03 | 0.06 | 2.58 |
| Absolute Kvert | 0.82 (0.51 0.93) | 0.81 (0.49 0.93) | 2.91 | 5.71 | 11.41 | 0.85 (0.67 0.94) | 0.85 (0.66 0.94) | 3.68 | 7.22 | 11.68 |
| Normalized Kvert | 0.79 (0.42 0.92) | 0.78 (0.40 0.92) | 46.04 | 90.25 | 11.31 | 0.89 (0.75 0.95) | 0.89 (0.75 0.95) | 47.98 | 94.04 | 11.65 |
| PPP | 0.69 (0.12 0.89) | 0.68 (0.11 0.88) | 6.43 | 12.60 | 10.18 | 0.97 (0.93 0.98) | 0.97 (0.93 0.98) | 1.96 | 3.84 | 6.17 |
| VTTS | 0.51 (0.00 0.82) | 0.50 (0.00 0.82) | 0.08 | 0.15 | 5.27 | 0.61 (0.15 0.84) | 0.61 (0.15 0.85) | 0.12 | 0.24 | 4.01 |

**Table 3.** Reliability analysis of outcome measures during double leg drop jump test

|  | Inter-day reliability metrics |  |  |  |  | Intra-day reliability metrics |  |  |  |  |
| --- | --- | --- | --- | --- | --- | --- | --- | --- | --- | --- |
|  | ICC(A,k) | ICC(C,k) | SEM | MMDC(±) | CV% | ICC(A,k) | ICC(C,k) | SEM | MMDC(±) | CV% |
| JH-TIA | 0.88 (0.66 0.95) | 0.87 (0.65 0.95) | 2.81 | 5.51 | 10.68 | 0.94 (0.87 0.98) | 0.95 (0.89 0.98) | 2.44 | 4.79 | 8.59 |
| JH-TOV | 0.88 (0.68 0.95) | 0.88 (0.67 0.96) | 2.78 | 5.44 | 11.15 | 0.93 (0.85 0.97) | 0.94 (0.86 0.98) | 2.73 | 5.36 | 9.41 |
| GCT | 0.82 (0.48 0.93) | 0.81 (0.47 0.93) | 0.03 | 0.05 | 6.88 | 0.90 (0.77 0.96) | 0.89 (0.76 0.96) | 0.03 | 0.06 | 6.15 |
| RSI | 0.93 (0.80 0.97) | 0.93 (0.80 0.97) | 0.10 | 0.20 | 10.17 | 0.94 (0.87 0.98) | 0.94 (0.87 0.98) | 0.11 | 0.21 | 11.05 |
| BPT | 0.82 (0.48 0.93) | 0.81 (0.47 0.93) | 0.01 | 0.03 | 7.23 | 0.87 (0.71 0.95) | 0.87 (0.70 0.95) | 0.02 | 0.03 | 7.69 |
| PPT | 0.81 (0.47 0.93) | 0.80 (0.46 0.93) | 0.01 | 0.03 | 6.96 | 0.91 (0.80 0.97) | 0.91 (0.79 0.96) | 0.01 | 0.03 | 5.63 |
| MBF | 0.88 (0.66 0.95) | 0.88 (0.66 0.96) | 0.17 | 0.33 | 5.13 | 0.93 (0.84 0.97) | 0.93 (0.83 0.97) | 0.18 | 0.35 | 5.04 |
| MPF | 0.92 (0.78 0.97) | 0.92 (0.77 0.97) | 0.14 | 0.27 | 4.33 | 0.93 (0.85 0.97) | 0.93 (0.85 0.97) | 0.15 | 0.30 | 4.91 |
| PGRF | 0.93 (0.79 0.97) | 0.92 (0.79 0.97) | 0.33 | 0.65 | 5.98 | 0.95 (0.88 0.98) | 0.95 (0.89 0.98) | 0.34 | 0.67 | 6.15 |
| PCOMD | 0.75 (0.30 0.91) | 0.74 (0.29 0.91) | 1.67 | 3.27 | 6.55 | 0.89 (0.76 0.96) | 0.90 (0.77 0.96) | 1.67 | 3.28 | 6.98 |
| RFD | 0.90 (0.71 0.96) | 0.89 (0.70 0.96) | 2.31 | 4.53 | 5.65 | 0.92 (0.82 0.97) | 0.92 (0.82 0.97) | 2.64 | 5.17 | 6.10 |
| VJI | 0.63 (0.00 0.86) | 0.62 (0.00 0.86) | 0.04 | 0.08 | 4.33 | 0.91 (0.80 0.96) | 0.91 (0.79 0.96) | 0.03 | 0.07 | 2.81 |
| Absolute Kvert | 0.61 (0.00 0.86) | 0.59 (0.00 0.85) | 5.87 | 11.50 | 14.55 | 0.93 (0.84 0.97) | 0.93 (0.84 0.97) | 3.88 | 7.61 | 12.04 |
| Normalized Kvert | 0.71 (0.18 0.89) | 0.70 (0.17 0.89) | 68.17 | 133.62 | 14.72 | 0.87 (0.72 0.95) | 0.87 (0.71 0.95) | 62.44 | 122.39 | 12.07 |
| PPP | 0.93 (0.82 0.97) | 0.93 (0.82 0.98) | 4.03 | 7.90 | 6.71 | 0.94 (0.87 0.98) | 0.94 (0.87 0.98) | 4.67 | 9.16 | 7.27 |
| VTTS | 0.65 (0.07 0.87) | 0.66 (0.06 0.88) | 0.06 | 0.13 | 3.89 | 0.77 (0.47 0.91) | 0.76 (0.45 0.90) | 0.05 | 0.10 | 2.99 |

ICC (A,k): absolute agreement ICC, ICC (C,k): consistency ICC, Lower and upper bounds of 95% confidence intervals in parenthesis, SEM: standard error of measurement, MMDC: minimal metrically detectable change, CV: coefficient of variation, JH-TIA (cm): jump height by flight time in the, JH-TOV(cm): jump height from take-off velocity, GCT(s): ground contact time, RSI (cm/s): reactive strength index, BPT(s): braking phase time, PPT(s): propulsion phase time, MBF (N/kg): mean braking force, MPF (N/kg): mean propulsion force, PGRF (N/kg): peak maximal ground reaction force, PCOMD (cm): peak center of mass displacement, RFD (N/kg/s): rate force development, VJI (BW\*s): vertical jump impulse, Absolute Kvert (kN/m): absolute vertical stiffness, Normalized Kvert (N/kg/m): normalized vertical stiffness, PPP (W/kg): peak propulsion power, VTTS (s): vertical time to stabilization

Based on paired t-test results, there were no significant differences between the test and retest mean values for any outcome measures of single and double leg DJ tests (p > 0.05 for every comparison). This result would demonstrate absence of any systematic bias in the DJ tests from measurement to measurement.

For single leg DJ tests (Table 2), the inter-day ICC values of no outcome measures could be considered very high level of reliability according to Munro’s classification. The inter-day ICC values of the ten outcome measures, i.e., JH-TIA, JH-TOV, GCT, BPT, PPT, MPF, PCOMD, VJI, absolute Kvert, and normalized Kvert, ranged from 0.70 to 0.89, which could be considered high level of reliability. Also, the inter-day ICC values of five outcome measures, like RSI, MBF, RFD, PPP, and VTTS, ranged from 0.50 to 0.69, which could be considered moderate level of reliability. Lastly, PGRF was the only outcome measure that have low level of inter-day reliability. In terms of absolute reliability, the highest CV level was observed for VJI as 2.82%.

For double leg DJ tests (Table 3), the inter-day ICC values of the four outcome measures, namely, RSI, MPF, PGRF, and PPP, ranged from 0.90 to 1.00, which could be considered very high level of reliability according to Munro’s classification. Similarly, the inter-day ICC values of the nine outcome measures, i.e., JH-TIA, JH-TOV, GCT, BPT, PPT, MBF, PCOMD, RFD, and normalized Kvert, ranged from 0.70 to 0.89, which could be considered high level of reliability. Lastly, the inter-day ICC values of only three outcome measures, like VJI, absolute Kvert, and VTTS, ranged from 0.50 to 0.69, which could be considered moderate level of reliability. On the contrary, absolute reliability was the highest for VTTS based on CV level of 3.89%.

In terms of intra-day ICC and double leg condition (Table 3), very high reliability was observed for RSI with ICC value of 0.94 and CV level of 11.04%. Similarly, for single leg DJ test (Table 2), very high reliability was evidenced for RSI with ICC value of 0.97 and CV level of 14.10%. Among all intra-day ICC values, the smallest value was demonstrated for VTTS with ICC value of 0.61 in the single leg DJ test.

## Discussion

This reliability study was conducted for presenting reliability metrics of outcome measures of single and double leg DJ tests. For the double leg DJ test, the results of the study indicated that except VJI, absolute Kvert, and VTTS, the inter-day ICC values of all the other outcome measures computed in this study could be considered to have high to very high level of reliability according to Munro’s classification (Carter and Lubinsky, 2016). On the other hand, for single leg DJ test, inter-day ICC values of ten outcome measures could be considered to have high level reliability. The reliability of jump height measure obtained by two methods, namely, flight time in the air (JH-TIA) and the vertical velocity of BCOM at the take-off instant (JH-TOV) were virtually the same in terms of the absolute agreement and consistency ICC values. Actually, near equality of absolute agreement and consistency ICC values was present for all outcome measures computed in this study which would indicate absence of bias or systematic error in the DJ measurements with a portable force plate (Liljequist et al., 2019).

In general, the inter-day ICC values of outcome variables in the double leg DJ test were higher than the corresponding ones in the single leg DJ test (11 out of 16). Conversely, the intra-day ICC values of outcome variables in the single leg DJ test were greater than the corresponding ones in the double leg DJ test (12 out of 16). Baxter et al. (2021) suggested that single and double leg DJs are nonidentical exercises in terms of Achilles tendon loading profiles. That would imply that single and double leg DJs could be evaluated separately in terms of SSC function as loading profiles are different. The result of this study presented that inter-day and intra-day ICC values of outcome measures in single and double leg DJs would be different and it should not be assumed to be always superior for the double leg DJ than the single leg DJ which is more challenging in terms of Achilles tendon loading.

The inter-day and intra-day ICC values of JH calculated by two methods using the flight time in the air (JH-TIA) and the vertical velocity of BCOM at the take-off instant (JH-TOV) were almost identical. When a jump mat is used for measuring jumping performance, the flight time method is used to calculate JH. However, when a force plate is used, either method could be employed in the calculations. Moir (2008) suggested that both methods are valid and consistent, however, when a force platform is available, practitioners or experimenters should use the TOV method for countermovement jump. We wanted to see whether there is a large difference in terms of reliability metrics hence it would be possible to favor one method to the other for the DJ tests, yet, the results of the reliability analysis yielded very similar values for two methods. Nevertheless, the CV values were slightly lower for the JH-TIA, therefore, time in the air method might be favored for a little better absolute reliability.

Among the outcome measures of single and double leg DJ tests, VTTS had the lowest or the second lowest inter-day and intraday ICC values. VTTS, or time to stabilization measures for any direction have been offered to analyze dynamic postural stability (Ross and Guskiewicz, 2008). Assessing dynamic stability using dynamic tests such as single and double leg DJ would especially be critical to detect postural instabilities in athletes (Ross and Guskiewicz, 2008). For that purpose, many methods had been developed to estimate time to stabilization measures (Fransz et al., 2015). Previously, Flanagan et al. (2008) assessed reliability of VTSS using a method in which VTSS was located as the first instance when VGRF reached and stayed within a threshold for one second, and the analysis yielded an intra-day ICC value that could be considered as moderate level of reliability. To estimate VTTS, we used sequential estimation method which was shown to be able to identify postural instabilities (Colby et al., 1999). For sequential estimation method, the results of current study with collegiate athletes demonstrated ICC values which could be considered moderate to high level of reliability and rather low CV values (CV < 10%).

Certain limitations affected our study. In this study, we studied 16 outcome measures, yet there is possibility to extract more information from DJ test, an instrumental jump test which has the potential to generate reliable and rich data sets to study biomechanical performance, neuromuscular control, and SSC function. Future studies may consider adding more outcome measures, possibly combining data from other sources as electromyography, three-dimensional motion capture, and modeling and simulation. In addition, it is possible that some choices on detecting key-time points might affect the reliability and magnitude of outcome measures. A recent study showed that changing threshold values to detect take-off instance influences reliability and magnitude of countermovement jump outcome measures (Perez-Castilla et al., 2021). Additional reliability studies may study the same issue for the DJ test.

In conclusion, the suggested DJ outcome measures (Bishop et al., 2021) as RSI and its individual components JH and GCT as well as normalized Kvert yielded high inter-day and intra-day ICC and low SEM and CV levels for the double leg DJ test. Those results thus suggest that the double leg DJ test could be used to assess jumping performance by using those above-mentioned outcome measures in collegiate handball players. The more challenging single leg DJ test could also be used in assessments with some other four reliable outcome measures like JH, GCT, normalized Kvert, and VJI. In conclusion, DJ tests could be deemed reliable to be used for short-term and long-term monitoring from the point view of biomechanical performance, neuromuscular control, and SSC function.

## Acknowledgments

This research was partially supported by The Scientific and Technological Research Council of Turkey under Grant 219S813.

